# Dynamic Multiday Seizure Cycles and Evolving Rhythms in a Tetanus Toxin Rat Model of Epilepsy

**DOI:** 10.1101/2024.10.15.618613

**Authors:** Parvin Zarei Eskikand, Mark J. Cook, Anthony N. Burkitt, David B. Grayden

## Abstract

Epilepsy is characterized by recurrent, unpredictable seizures that impose significant challenges in daily management and treatment. One emerging area of interest is the identification of seizure cycles, including multiday patterns, which may offer insights into seizure prediction and treatment optimization. This study investigated multiday seizure cycles in a Tetanus Toxin (TT) rat model of epilepsy. Six TT-injected rats were observed over a 40-day period, with continuous EEG monitoring to record seizure events. Wavelet transform analysis revealed significant multiday cycles in seizure occurrences, with periods ranging from 4 to 7 days across different rats. Synchronization Index (SI) analysis demonstrated variable phase locking, with some rats showing strong synchronization of seizures with specific phases of the cycle. Importantly, the study revealed that these seizure cycles are dynamic and evolve over time, with some rats exhibiting shifts in cycle periods during the recording period. This suggests that the underlying neural mechanisms driving these cycles may change as the epileptic state progresses. The identification of stable and evolving multiday rhythms in seizure activity, independent of external factors, highlights a potential intrinsic biological basis for seizure timing. These findings offer promising avenues for improving seizure forecasting and designing personalized, timing-based therapeutic interventions in epilepsy. Future research should explore the underlying neural mechanisms and clinical applications of multiday seizure cycles.

## Introduction

Epilepsy, a chronic neurological disorder characterized by recurrent seizures, affects millions worldwide, presenting a significant burden on individuals and healthcare systems. Despite advancements in treatment, many patients continue to experience unpredictable seizure occurrences, highlighting the complex and heterogeneous nature of the disease. One of the most challenging aspects of epilepsy is the apparent randomness of seizure occurrences, which significantly complicates the lives of patients. Understanding and studying the temporal patterns and cycles in epilepsy can provide valuable insights into seizure forecasting and also the underlying mechanisms of seizure generation and progression.

Multi-day cycles in epilepsy refer to recurrent patterns in seizure occurrence over extended periods, often displaying rhythmic variations in frequency and intensity. Understanding these patterns is crucial as they hold potential implications for seizure prediction, treatment optimization, and ultimately, improving patient outcomes. By deciphering the temporal dynamics of epilepsy through multi-cycle analysis, researchers aim to uncover predictive biomarkers, refine therapeutic strategies tailored to individualized seizure patterns, and advance our understanding of the disease’s pathophysiology. Recent research has increasingly recognized the importance of temporal patterns and cycles in epilepsy, beyond mere sporadic events [19, 20]. Over the past decade, the analysis of extensive EEG datasets has advanced significantly, revealing substantial evidence that epilepsy cycles occur at various timescales—such as circadian, multidien, and circannual. These cyclical patterns are widely observed in both human and animal studies of epilepsy, with evidence from sources like implantable devices, electronic seizure diaries, and wearable monitoring systems [1, 2, 7, 12, 17, 18, 28].

In this study, we focus on the TT rat model of epilepsy to investigate the presence and characteristics of multiday cycles in seizure occurrences. The TT model, known for its robust and reproducible induction of chronic epilepsy, provides an excellent platform for studying the temporal dynamics of seizures. The TT rat model is commonly used to study temporal lobe epilepsy (TLE), which is one of the most common forms of epilepsy in humans. TLE is often characterized by complex partial seizures that can evolve over time, similar to the progression observed in the TT model. By analyzing long-term EEG recordings from this model, we aim to identify and characterize multiday cycles, providing insights into their potential mechanisms and implications for therapeutic interventions.

## Materials and methods

### Data

The data used in this study was obtained from a intra-hippocampal TT rat model of epilepsy [8]. Six adult male Sprague-Dawley rats, sourced from the Animal Resources Centre, were used in the study, with intra-hippocampal injections of TT administered to form the TT group, while four control rats received phosphate-buffered saline. On the day of surgery, their weights ranged between 250–370 g. The animals were housed individually in acrylic cages and provided with unrestricted access to food and water. Environmental conditions in the recording room were carefully maintained, including a controlled temperature range of 21–26°C. The light/dark cycle consisted of 16 hours of light (approximately 16:15–08:08 Australian Eastern Standard Time) followed by 8 hours of darkness (approximately 08:08–16:15 AEST), with the transitions managed by an automated timer.Prior to surgery, rats were anesthetized with Ketamine (75 mg/kg) and Xylazine (10 mg/kg). Each rat had five stainless steel electrodes implanted into their skulls to allow for intracranial EEG recording.Using the bregma as the origin at point (0,0), the electrode placements (anteroposterior, mediolateral) were positioned as follows (in mm): B (1.2, +3.0), W (6.8, +3.0), G (1.2, −3.0), R (6.8, −3.0), and Ref (10.6, −3.0). All other methods related to the collection of the data used in our study are contained in the original paper by Crisp et al. (2020) [8]. TT was administered stereotaxically in a volume of 300 nanoliters to rats in the Epilepsy Model Group, delivering a dose of 30 nanograms. In the Control Group, the same procedure was followed, but only vehicle solution was used. The injection was targeted to the CA3 region of the hippocampus, with stereotaxic coordinates 3.5 mm posterior to bregma, 3.0 mm lateral to the midline (left hemisphere), and 3.1 mm below the dura mater. Approximately 1-2 weeks after the TT injection, the TT rats began experiencing spontaneous seizures.

The detection of spontaneous seizures was performed using EEG recordings analyzed with a custom-designed algorithm. EEG signals were processed using a high-pass filter (with a cutoff frequency at 1 Hz) to remove noise and artifacts. The power of the EEG signal was computed within the 13–48 Hz frequency band. This range was chosen based on its relevance in detecting epileptiform activity associated with seizures. A potential seizure was flagged if the instantaneous band power exceeded 3 times the 5-minute moving average for at least 9 seconds or 7 times for at least 3 seconds. These thresholds were carefully selected to balance sensitivity and specificity in identifying epileptic events. For a seizure event to be confirmed, the potential seizure needed to be simultaneously detected on at least two electrodes. This criterion minimized false positives due to localized noise or artifacts. The onset of a seizure was defined as the time when the band power first rose above the threshold. The termination of a seizure was marked when the band power returned to or below the moving average. These rigorous criteria ensured the reliable identification of spontaneous seizures in the EEG data [5, 8]. The EEG trace in Fig. S1 illustrates a representative spontaneous seizure observed in the epilepsy model group. The spontaneous seizure studied in this work are electrographic seizures without the behavioral component.

The number of daily seizures fluctuated over a span of about six weeks, as shown in Fig. 1. Around weeks 4 to 5 post-injection, the seizure frequency started to decline, eventually resulting in no detectable seizures. EEG data was continuously recorded at a sampling rate of 2048 Hz for 23 hours each day, with 1 hour allocated for daily maintenance and data backup. Of the original seven TT rats, six completed the study, as one died on Day 26.

**Fig 1.**
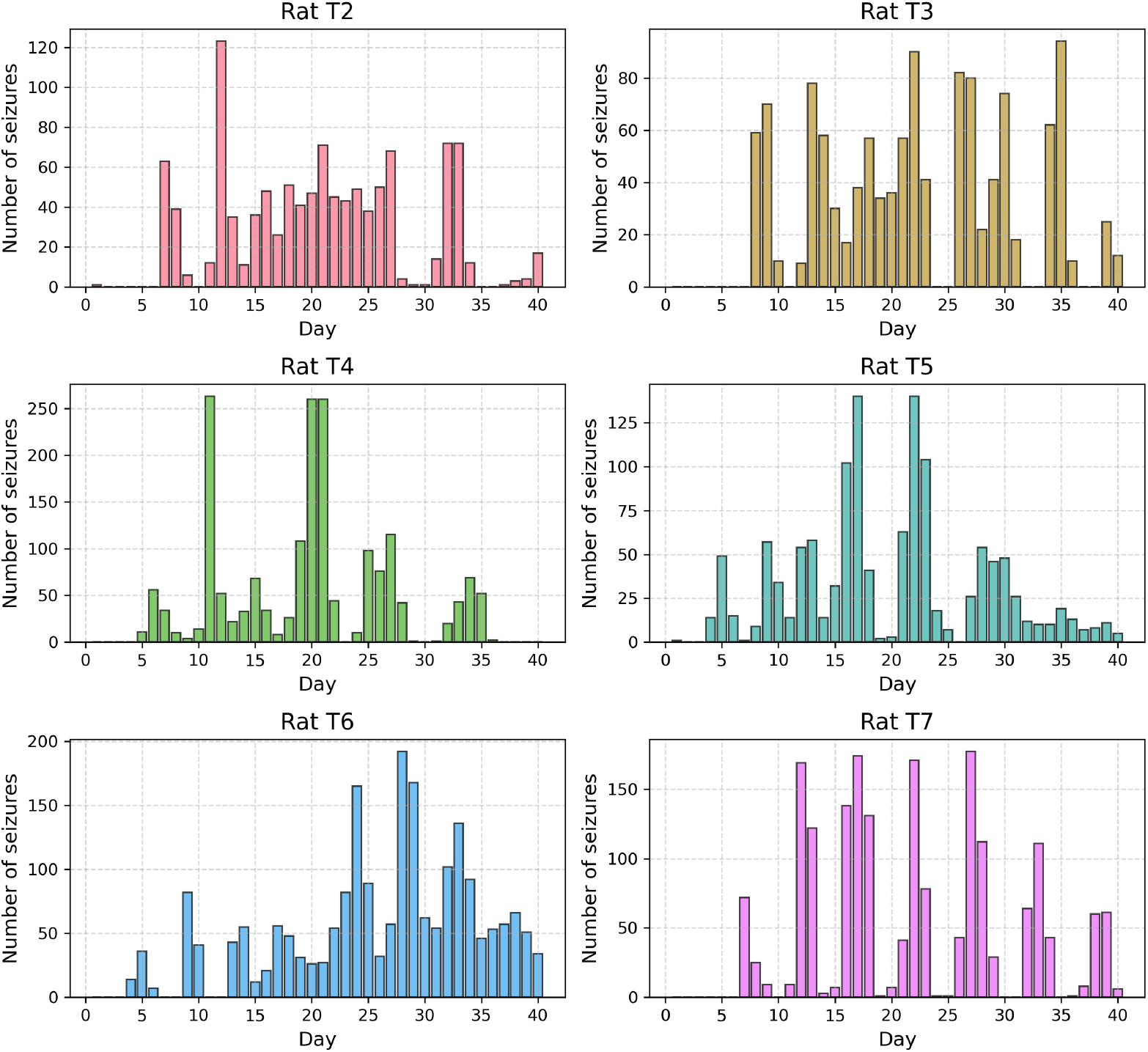
Seizure occurrences over time for TT rats. Each subplot represents data from one rat with the x-axis indicating days since the start of the recording. The y-axis is the total number of seizures that occurred on the specific day.

Fig. S2 shows the distribution of seizure frequencies per hour over a period of 40 days for six rats. Some hours of the day show a drop in the number of seizures, and this pattern varies between different rats. Fig. 2 shows the number of seizures recorded on specific days from day 1 to day 40, as well as at particular hours of the day. The seizure durations varied significantly among the six rats (T2–T7), with each displaying a unique range [32][2]. The study by [32] identified significant cyclical patterns in seizure durations over time, with periods ranging from 4.1 to 8.6 days across the rats. These cycles capture the rhythmic fluctuations in seizure lengths, which were not static but evolved during the 40-day recording period [32].

**Fig 2.**
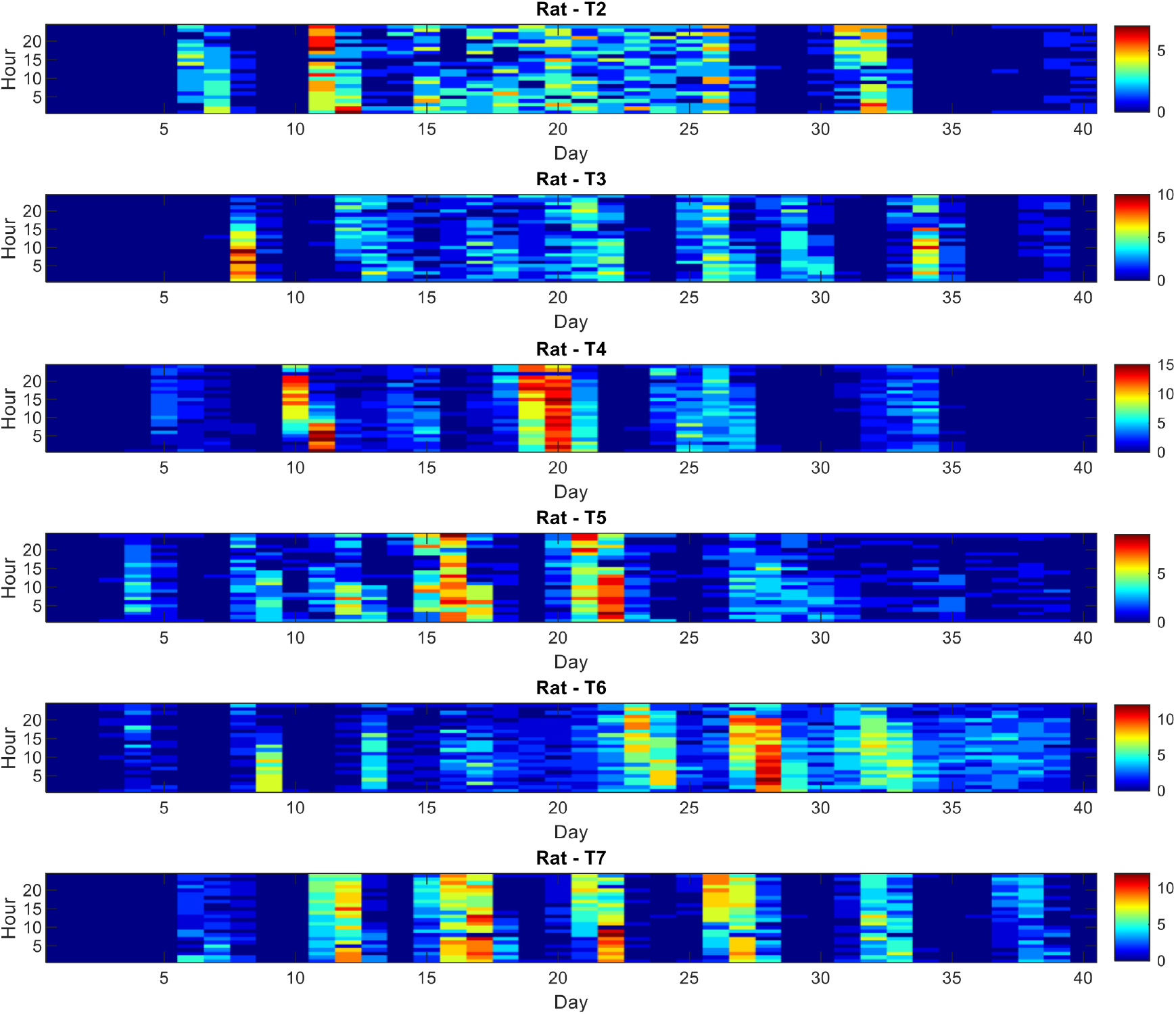
Seizure distribution across days and hours for six TT rats. The distribution of seizures recorded over a period of 40 days, plotted against hours of the day. The x-axis represents the day of recording, while the y-axis denotes the hour of the recording session. Colors ranging from red to blue indicate varying frequencies of seizures, with red indicating higher frequencies and blue indicating lower frequencies.

The experimental design involved periodic low-level electrical stimulation, referred to as “probing” [8]. Each probing-on phase consisted of delivering 100 biphasic pulses, with a pulse width of 0.5 ms and a current of 1.26 *±* 0.024 mA per phase, spaced by inter-stimulus intervals of 3.01 s, lasting for approximately 5 minutes. This was followed by a 5 minute probing-off phase before the next probing-on session began. Although probing was part of the experimental protocol, it did not play a role in the specific analyses or outcomes investigated in this study. In our study, control rats received identical surgical and handling procedures as the epilepsy model group, except that they were injected with saline without TT. These animals exhibited no spontaneous seizures throughout the study, and their EEG recordings served as baseline data to distinguish pathological activity related to epilepsy. The study was conducted in compliance with ethical standards, receiving approval from the St. Vincent’s Hospital Melbourne Animal Ethics Committee, in accordance with the “Australian Code for the Care and Use of Animals for Scientific Purposes, 8th Edition” (2013). This study is reported in accordance with the ARRIVE guidelines (Animal Research: Reporting of In Vivo Experiments) to ensure comprehensive and transparent reporting of the research methods and findings [25].The data analyzed in this study were previously obtained (Crisp et al., 2020 and Cheung, 2017) [8].

### Temporal analysis of seizure occurrences

The number of daily seizures occurring per hour over a 40-day recording period for six rats was collected. This data was analyzed to identify underlying periodic patterns using Wavelet transforms and sinusoidal curve fitting [20]. We utilized the Continuous Wavelet Transform (CWT) to analyze the temporal patterns and multi-day cycles in seizure occurrences recorded over a span of 40 days. The dataset comprised hourly seizure counts across multiple rats, each represented as a time series. The CWT was applied using a Morlet wavelet, a well-suited wavelet for analyzing time series with transient signals [20]. This wavelet transformation facilitated the decomposition of the seizure count time series into its frequency components, revealing both localized and global power spectra. To identify significant frequency components corresponding to multi-day cycles, a range of frequencies spanning from 4 hours cycles per day up to 240 hours (10 days) cycles was explored. We set the maximum to 10 days to be able to observe at least four cycles of the data over 40 days.

Significance testing was performed to distinguish meaningful frequency components from noise. Significant peaks in the global wavelet power were determined using a time-averaged significance test as described in [30]. This is done by comparing the observed wavelet power against the distribution of power expected under the null hypothesis that the data is generated by the chosen background noise model (in this case, a red-noise process), with the significance level at 95% to account for autocorrelation properties and multiple comparisons [20]. Peaks exceeding the 95% significance level in the wavelet power spectrum were identified as significant cycles. These peaks corresponded to periods where seizure occurrences exhibited pronounced cyclical behavior across the 40-day observation period. The Synchronization Index (SI) was calculated for the significant peaks with the highest power. The SI index measures the phase consistency of seizures within the data. This was done by converting the timestamps of seizures into phase values and computing the mean resultant vector [20]. The SI index, *I*_*S*_, is a measure of the phase locking of seizure occurrences, with values ranging from 0 to 1. Higher SI values indicate stronger synchronization in seizure timings. This calculated using the following equation:

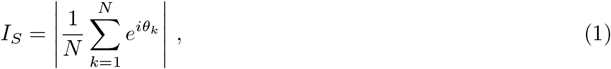

where *N* is the number of events, *θ*_*k*_ is the phase of the *k*-th seizure, and 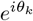 represents the complex exponential of the phase. The SI is the magnitude of the mean resultant vector, calculated by taking the absolute value of the average of these complex exponentials. The SI value ranges from 0 to 1, where 0 indicates no synchronization and 1 indicates perfect synchronization [20].

For better visualization of the cycles, we also fitted a sinusoid to the hourly number of seizures for each rat. The sinusoidal function used is defined as:

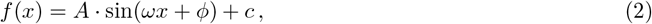

where *A* is the amplitude, *ω* is the angular frequency, *ϕ* is the phase shift, and *c* is the offset. The optimization was performed by minimizing the Sum of Squared Residuals (SSR) between the observed data and the sinusoidal model. The periods identified for each rat were recorded and reported.

As the final step, to better illustrate the effect of TT on the cycles, the data was segmented into 15-day intervals (360 hours each) with 359 hours overlap to analyze periodic patterns over time. For each segment, the significant peaks in the cycles were calculated using the method described above.

We also calculated the significance of the peak by comparing the observed power at that frequency to the global significance threshold obtained from the wavelet power spectrum.

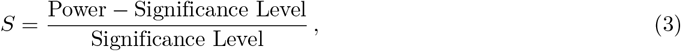

Where Power is the observed power at the specific frequency and Significance Level is the threshold power level determined by the global significance test. Higher *S* values indicate greater significance, with the color transitioning from cooler tones (e.g., blue) for lower *S* values to warmer tones (e.g., purple) for higher *S* values. This allows for a visual representation of the relative strength of each significant peak in the frequency domain.

## Results

The analysis of seizure data for six rats over a period of 40 days revealed distinct periodic patterns for each rat. Fig. 3 illustrates the wavelet power spectrum analysis results for each rat. The power spectrum depicts the distribution of power across different periods (in hours), highlighting significant peaks where seizure occurrences exhibited pronounced cyclical behavior over the 40-day observation period. Significant peaks were determined using a time-averaged significance test (*α* = 0.05), accounting for autocorrelation properties and multiple comparisons. Significant peaks in the global wavelet power spectrum were identified as periods where the period of seizure occurrences were statistically higher than expected by chance. These peaks, marked in the figure as ‘Significant Peaks’, indicate intervals of heightened seizure activity that recur periodically within each rat’s data. Peaks exceeding the 95% significance level (black dashed line) were considered robust indicators of cyclic behavior in seizure occurrence. Fig. 3 shows the presence of dominant multiple cycles in the seizure occurrence in the TT rat model of epilepsy. The cycles found for different rats are listed as follows: Rat T2: 157 hours (6.5 days), 121 hours (5 days), 73 hours (3 days); Rat T3: 159 hours (6.6 days), 104 hours (4.3 days); Rat T4: 174 hours (7.2 days); Rat T5: 145 hours (6 days); Rat T6: 114 hours (4.7 days); Rat T7: 126 hours (5.2 days).

**Fig 3.**
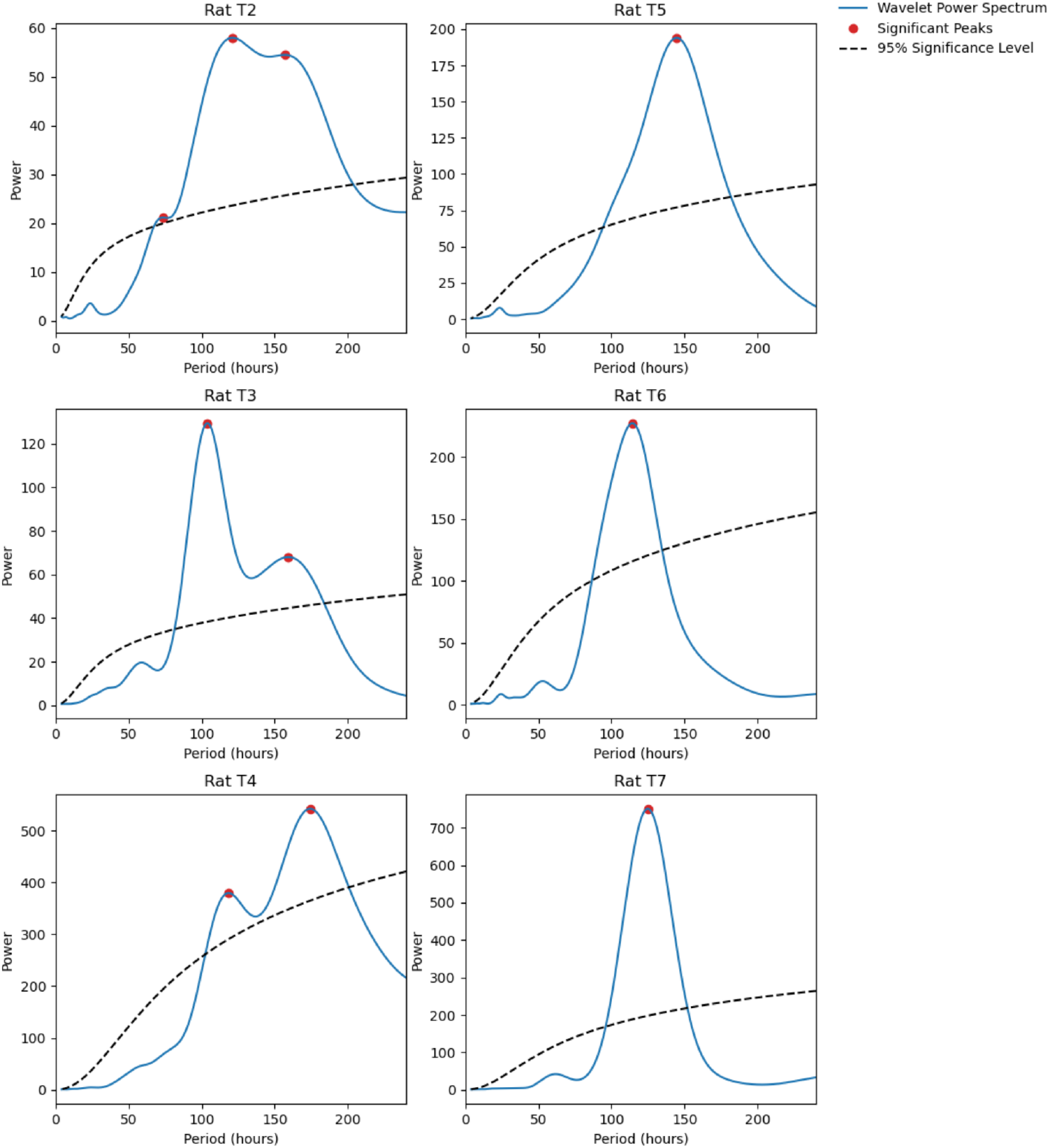
Wavelet power spectrum analysis of hourly seizure occurrences for seizure data recorded over 40 days. Red dots indicate significant peaks in the wavelet power spectra of numbers of seizures occurring per hour, which are indicated by the blue lines showing power at different periods (hours). Each subplot corresponds to a different rat (T2 to T7). The dashed black line represents the 95% significance threshold.

Using sinusoidal curve fitting, we identified the optimal parameters that best describe the cyclic nature of the seizure occurrences, assuming that cycle frequencies were fixed over the whole recording periods. Fig. 4 illustrates the fitted sinusoidal models and the corresponding numbers of seizures per hour for each rat. The observed data points are marked with blue stars, while the fitted curves are depicted in orange. The period of the sinusoidal function is annotated in each subplot, indicating the duration of the cycles in hours (days). The periods identified for each rat are consistent with the results obtained from the wavelet transform analysis that showed the strongest power. This figure better visualizes one of the rhythmic cycles identified in the data. In the following section, the temporal changes in these significant cycles over 40 days of recording are investigated.

**Fig 4.**
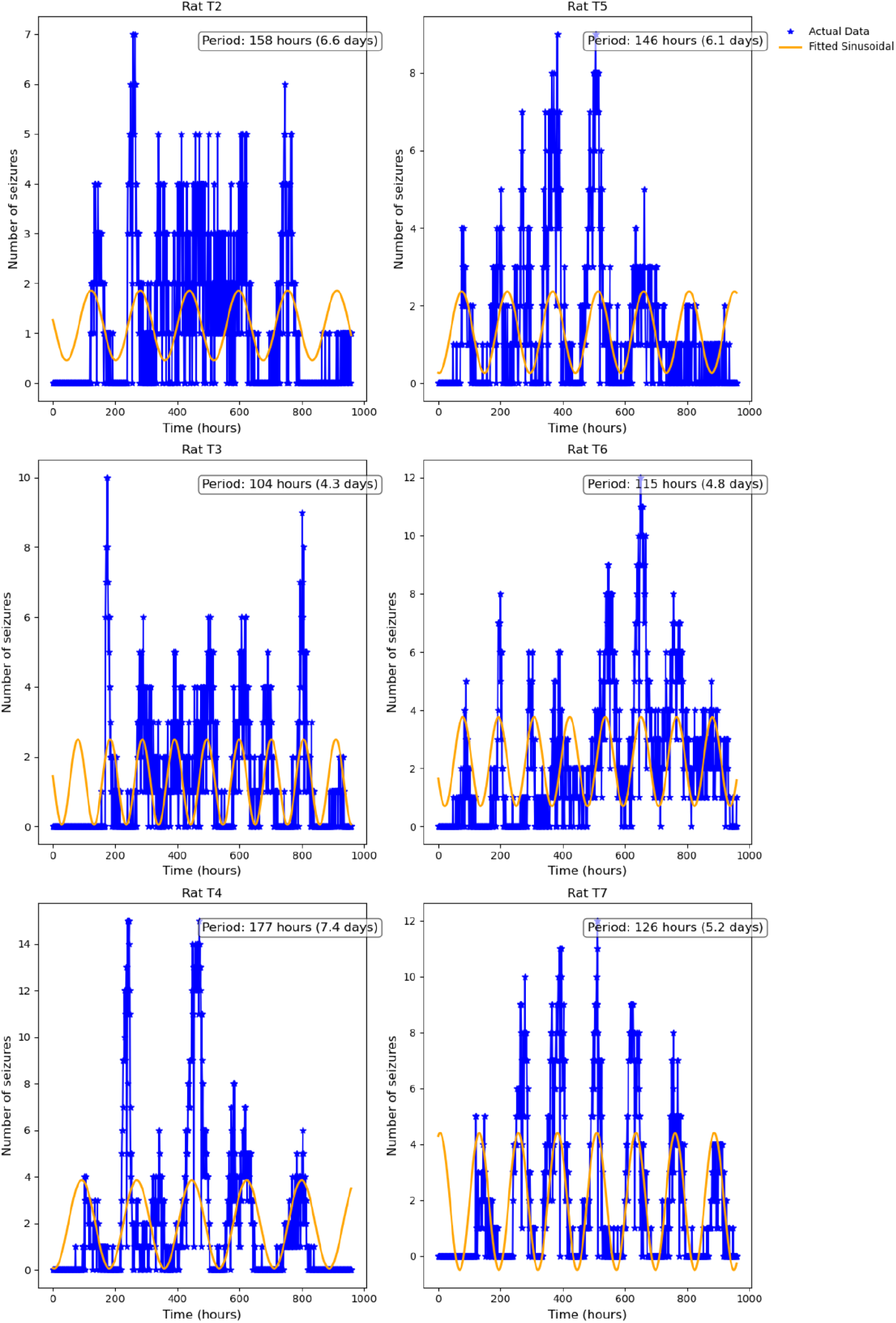
Sinusoidal curve fitting of seizure data. Each subplot displays the actual seizure counts (blue stars) and the fitted sinusoidal model (orange line). The period of the sinusoidal fit is annotated for each subplot, demonstrating the periodic patterns observed in seizure occurrences over time.

To further understand the cyclic nature of seizures in the rats, polar plots were generated to illustrate the distribution of seizure phases within one cycle of the most significant rhythmic period as determined by the wavelet transform analysis. Fig. 5 presents the polar histograms of seizure timings for each rat, based on the periods identified from the wavelet power spectrum. These polar plots help illustrate the phase locking of seizures and provide insights into the SI of seizure occurrences. The histograms are displayed in polar coordinates, where angles represent the phases of the cycle (processing clockwise) and the radial axis indicates the relative number of of seizure occurrences within each phase interval. For example, the polar histogram for Rat T7 demonstrates a strong clustering of seizures around the “Rising” phase, with very few events occurring during the “Falling” and “Trough” phases. The clustering of seizures in a specific phase interval implies that seizure occurrences are not random but follow a predictable pattern. For example, rat T7 has the highest SI of 0.64, indicating the strongest phase locking of all the rats to their dominant frequencies. This strong synchronization suggests that seizures in Rat T7 are highly predictable throughout the recording and occur predominantly during the “Rising” phase. This predictability could be useful for anticipating seizure events and implementing preventative measures.

**Fig 5.**
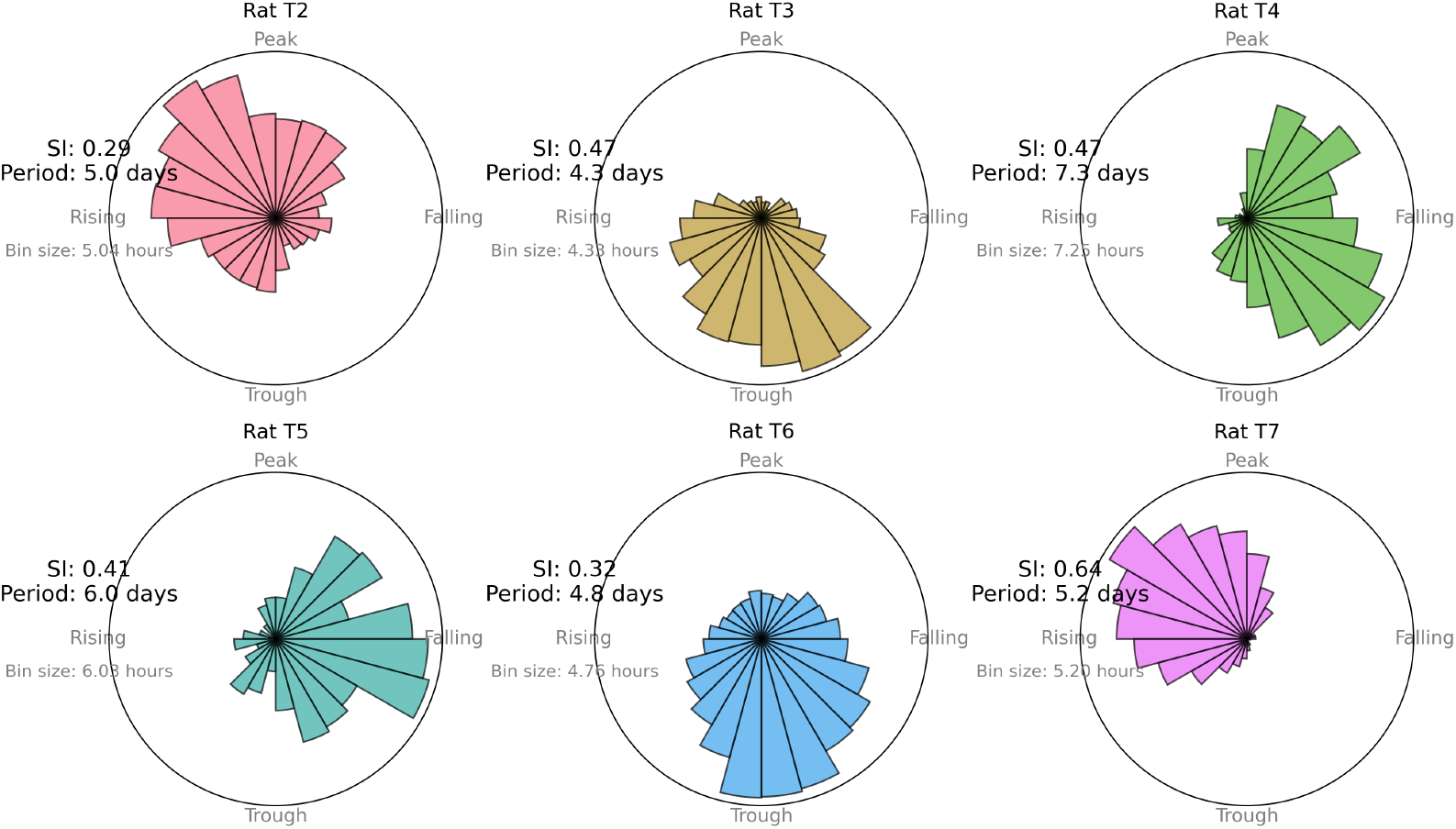
Polar histograms of seizure timings for each rat based on the identified rhythmic periods. Each subplot represents a different rat (T2 through T7). The radial axes indicate the frequency of seizure occurrences within each phase interval, and the angular axes represent the phases of the identified rhythmic cycle. The predominant period and the Synchronization Index (SI), Eq.(1), are annotated on each plot. Higher SI values indicate stronger phase locking and synchronization of seizure occurrences.

The TT model of epilepsy is known to wax and wane over the weeks of recording, and for the properties of the seizures to change over time [8]. To investigate these effects over time, the data was divided into segments, each 360 hours (15 days) long and advancing 1 hour between consecutive segments. This segmentation approach allowed for the analysis of seizure periods across different time windows along the 40 days of recording. For each segment, the wavelet power spectrum was computed. Peaks were considered significant if their power exceeded the threshold set by the significance level (0.95).

Fig. 6 illustrates the significant periods of seizures over the duration of recording. Each subplot represents one rat and displays colored dots corresponding to significant seizure periods over time (days) for multiple segments. The color of each line represents the Peak Significance *S* value, Eq. (3), shown on a log scale. Higher *S* values indicate greater significance, with the color transitioning from cooler tones (i.e., blue) for lower *S* values to warmer tones (i.e., purple) for higher *S* values. This allows for a visual representation of the relative strength of each significant peak in the frequency domain. Each data point shows the significant cycle(s) determined from the preceeding 360 hours (15 days), as indicated by the dashed lines on the zero y-axes for the first data point.

**Fig 6.**
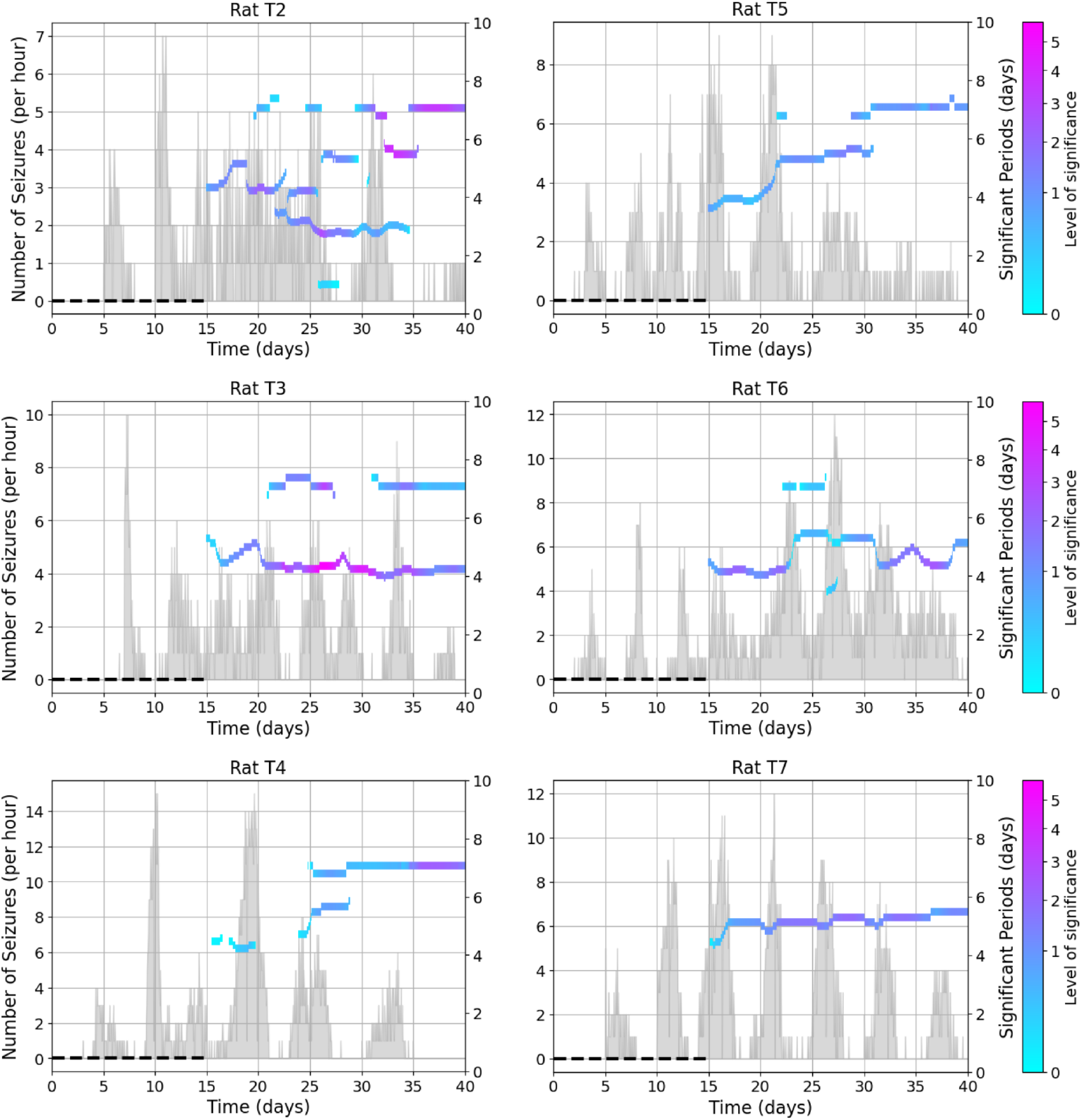
Significant cycles over time for different rats. Each subplot displays the significant periods (days) of seizures as horizontal lines, plotted against time (days) across multiple segments. Each segment lasts 360 hours (15 days) and overlaps with the next segment by 359 hours. The color of each line represents the significance of the peak, with the color bar on the right of each subplot indicating the scale, where lower significance values are shown in blue and higher significance values in purple. The gray histogram on the left axis shows the number of seizures per hour over the 40 days of recording. The dashed line indicated that the power of the wavelet calculated over a 15-day period.

This analysis presents the evolution of significant cycles in seizure activity over 40 days. The results indicate that the significant cycles are not static but change over time, reflecting dynamic rhythmic patterns in seizure occurrences. Some cycles persist across multiple segments, while others appear or disappear as the recording progresses. This variation suggests that the underlying neural mechanisms generating these cycles may shift over time.

For Rat T2, the significant cycles show considerable variation throughout the 40-day period. Early in the recording, cycles of around 4-5 days are consistently present, but later segments reveal the emergence of longer cycles (up to 7 days). There is notable variation in the significant periods over time, with some periods disappearing and reappearing. The significance of the power of some periods increases towards the end of the recording, with some high-power cycles (depicted in purple) emerging after 32 and 36 days. The results suggest that the rhythmic periods of seizures in Rat T2 evolve over time, with certain cycles becoming more pronounced (higher power) later in the recording. This indicates possible changes in the underlying mechanisms driving seizures as the experiment progresses.

Initially, Rat T3 shows significant cycles with periods around 4-5 days (approximately 100-120 hours) during the first 20 days of the recording. However, after day 20, a new dominant cycle with a period of around 7 days (approximately 170 hours) emerged. This 7-day cycle did not remain stable — it disappeared and reappeared intermittently until the end of the recording. This suggests that, while the 4-5-day cycles are consistently present throughout, the 7-day cycle was more transient, appearing sporadically. This variability could reflect changes in the underlying neural activity influenced by the progression of the TT effect or other factors.

Rat T4 shows more variability in the significant cycles. Early in the recording, cycles were around 4 days, but mid-recording, there was more variation, with cycles ranging from 4-5 days. Towards the end of the recording period, the cycles tended to stabilize around 7 days. This variability indicates a more complex seizure pattern with an increase over time.

In Rat T5, the significant cycles started with a period of around 4 days and gradually increased to around 7 days by the end of the recording. Around day 30, there was a period where two dominant cycles coexisted—one around 5.5 days and the other around 7 days. This indicates a shift in the rhythmic patterns of seizure occurrences.

Rat T6 showed a gradual change in the significant periods over the 40 days of recording. The significant cycles began with a period of around 4 days and then fluctuated, gradually increasing and decreasing over time. Between days 23-27, there were two dominant cycles present simultaneously, one shorter and one longer, indicating a temporary coexistence of different rhythmic patterns.

Rat T7 showed very stable significant cycles, maintaining a cycle length of around 5 days (120-130 hours) throughout the entire 40-day period. The cycles remained stable, indicating a strong rhythmic pattern in seizure occurrence. The cycles in Rat T7 were characterized by relatively high significance, reflecting strong synchronization of seizure activity.

## Discussion

TT model induces spontaneous seizures through a specific pathophysiological mechanism involving the disruption of inhibitory neurotransmission. TT is a neurotoxin that selectively targets inhibitory interneurons in the central nervous system. It enters presynaptic terminals of these neurons and cleaves synaptobrevin (also known as vesicle-associated membrane protein or VAMP) [15, 24].

By cleaving synaptobrevin, TT effectively blocks the release of inhibitory neurotransmitters such as inhibitory amino acid transmitters (GABA) and glycine. This blockade leads to a reduction in inhibitory synaptic transmission, creating an imbalance between excitatory and inhibitory inputs in neural networks. The decreased inhibitory tone results in neuronal hyperexcitability, promoting the generation and propagation of spontaneous epileptiform discharges and seizures [24].

Over time, the effects of TT diminish as the toxin is metabolized and synaptic proteins are resynthesized, gradually restoring normal inhibitory neurotransmission. This recovery process re-establishes the balance between excitation and inhibition in the brain, leading to a decrease and eventual cessation of spontaneous seizures [16].

In this study, we examined the rhythmic patterns of seizure occurrences in a TT rat model of epilepsy using wavelet transform analysis to identify significant cycles in seizure timing. By analyzing the seizure data recorded over 40 days across six different rats (Rats T2-T7), we uncovered distinct rhythmic patterns that varied in both period and power across individual rats. These findings provide new insights into the complex dynamics of seizure occurrence and the impact of cyclic rhythms in epilepsy.

### Wavelet power spectrum analysis

The wavelet power spectrum analysis, shown in Fig. 3, provides a detailed view of the power and significance of different rhythmic periods in the seizure data. Across the six rats, we observed significant peaks corresponding to multi-day cycles, with periods typically ranging from 73-174 hours. For example, the presence of a single dominant peak at 120 hours suggests that Rat T7’s seizures were strongly synchronized with a 5-day cycle. The lack of additional significant peaks indicates that this period was the primary driver of rhythmic activity in Rat T7’s seizures, making it a highly consistent and predictable pattern. Some rats, like T5, T6 and T7, displayed one very strong and dominant cycle, while others, like T4, T2 and T3, showed multiple significant periods, indicating more complex seizure patterns. These multiple periods may correspond to different underlying processes that contribute to seizure generation, potentially offering multiple targets for therapeutic intervention. These findings suggest that the seizures in each rat are influenced by specific, recurring rhythms, which vary in strength and periodicity across different individuals.

### Temporal dynamics of significant rhythms

The analysis of significant periods over time, as depicted in Fig. 6, showed that the seizure rhythms were not static but evolved during the recording period. Some rats, like T3 and T7, showed stable rhythmic periods, while others, such as T2, exhibited more fluctuation and variability. Some rats, such as T2, T4 and T5, showed an increase in the period of significant cycles toward the end of the 40-days of recording. These findings suggest that the effects of TT on seizure dynamics may not be uniform across different rats. The observed shifts in rhythmic periods could reflect adaptive or pathological changes in the brain’s oscillatory networks, possibly related to the progression of the epileptic state or the brain’s response to either sustained seizure activity or the TT. Understanding the evolution of these significant seizure periods and their power can provide valuable insights into the effects of TT on neural rhythms and may inform the development of targeted interventions based on the specific rhythmic characteristics of each rat.

Crisp et al. (2020), also observed temporal changes in seizure dynamics in their study, showing that the type of seizure onset evolved over the recording period for these same rats. Initially, seizures were characterized by fast, low-amplitude spiking with a Direct Current(DC) shift at the beginning. However, as epileptogenesis progressed, seizures began to exhibit no DC shift, with repetitive bursts and increasing frequency toward the end. Additionally, changes in evoked potentials were noted, reflecting the brain’s increasing excitability in response to stimulation as the number of seizures increased [8]. It would be interesting to explore the relationship between the evolving seizure cycles in our study and the changes in seizure types and neural excitability observed by [8]. Specifically, investigating how the rhythmic periods of seizures may influence or be influenced by the underlying neural mechanisms driving these seizure onset dynamics could provide deeper insights into the effects of TT. This connection between the temporal evolution of seizure rhythms and seizure dynamics will be a focus of future studies.

### Importance of studying seizure cycles in animal models compared to humans

In humans, seizure cycles are often influenced by a multitude of external factors, which can complicate the analysis of their underlying physiological causes. For instance, differences in daily activities, such as altered routines during weekends, irregular sleep patterns [29], and increased stress or social interactions, can affect seizure timing [13, 14]. Environmental influences like air pollution [4] and diet [13] also play significant roles in seizure triggers and modulate the rhythms of seizure occurrences. These factors, while highly relevant to real-world human conditions, introduce considerable variability, making it difficult to isolate the core physiological mechanisms that generate seizure rhythms. By maintaining consistent lighting conditions, controlled feeding schedules, and a stable environment for the rats, we can effectively eliminate many of the external factors that complicate human studies.

Our study demonstrated that multi-day cycles in seizure occurrences persist even in the absence of the external factors typically observed in human studies, such as weekly changes in life routines, environmental pollution, and dietary variations. The presence of these multi-day rhythmic patterns in rats, despite the removal of such external factors, strongly suggests that these cycles have a physiological basis independent of environmental and lifestyle variables. This finding is particularly significant as it underscores the inherent nature of these seizure rhythms, pointing to underlying biological mechanisms—possibly related to circadian rhythms [20], endocrine systems [26], or other neurophysiological processes—that are responsible for generating these cycles. It highlights the importance of considering these intrinsic factors when studying epilepsy and designing interventions, as it indicates that the seizure cycles observed in humans are not solely the result of external influences, but are likely driven by deeper, endogenous processes. This insight provides a more solid foundation for understanding seizure dynamics and developing targeted therapeutic strategies that address the root causes of rhythmic seizure activity for each individual patient.

While the TT model provides robust and reproducible data for studying multiday seizure cycles, its applicability to human epilepsy requires further validation. Human epilepsy is influenced by a wide range of external factors, including environmental and behavioral changes, which were controlled in this study. While our findings demonstrate the presence of rhythmic seizure patterns, the underlying neural mechanisms driving these cycles remain speculative. Further studies combining computational modeling and experimental approaches are necessary to elucidate these mechanisms.

### Multi-days cycles in other rat models of epilepsy

There are other rat models of epilepsy that have been employed to investigate the cyclic nature of seizure occurrences and their underlying mechanisms. Notably, the kainic acid and pilocarpine models of TLE have provided valuable insights into the existence of circadian and multidien rhythms in epilepsy. Both of these models show multidien rhythms ranging from 2-7 days. However, these cycles were not as strong as the ones we observed in the TT model of epilepsy, particularly in the kainic acid model [2]. There are also other studies that have examined multi-day cycles in female rats in relation to their menstrual cycle, but not in male rats [9, 23].

The pilocarpine model, described by Lévesque et al. [21, 22, 27], offers valuable insights into mesial temporal lobe epilepsy (MTLE). This model highlights a latent period characterized by neural network reorganization, during which significant changes in interictal spike activity and high-frequency oscillations (HFOs) occur. These findings underscore the potential of specific biomarkers, such as HFOs and interictal spikes, to predict seizure onset and progression. Similarly, our study identifies temporal dynamics in seizure occurrences that may be linked to underlying processes of epileptogenesis. These dynamics could reflect intrinsic oscillations in neural excitability, akin to those observed in the pilocarpine model, where distinct patterns of spike activity in the hippocampus and entorhinal cortex evolve during the latent phase and stabilize in the chronic phase.

### Clinical implications and future directions

The findings of this study underscore the importance of understanding the temporal dynamics and rhythmic patterns of seizures in epilepsy. The identification of significant rhythmic periods and phase locking in seizure occurrences suggests that these cycles could be exploited for therapeutic purposes, such as timing interventions to coincide with periods of high seizure risk. Additionally, the variability in rhythmic patterns across different rats highlights the need for personalized approaches to epilepsy treatment, taking into account the specific rhythmic dynamics of each patient.

By identifying the specific cycles of seizure occurrence in individual patients, clinicians could tailor treatment schedules to align with periods of heightened seizure risk. For example, patients could receive prophylactic treatments or adjust their medications during periods when seizures are more likely to occur, thus improving seizure control while minimizing unnecessary side effects during low-risk periods. It has been demonstrated that responsive neurostimulation has significantly different effects when using standard parameters, depending on when stimulation occurs during the seizure cycle [6], which has important clinical consequences. Additionally, understanding these cycles could also help in predicting the likelihood of seizure clusters, providing patients with actionable information to prevent injury or seek timely medical intervention.

Detecting dynamic cycles of seizures in patients, particularly when these cycles may vary between individuals, can be achieved through wearable devices. These devices, capable of continuous and long-term monitoring, could record physiological signals such as EEG, heart rate, and movement patterns. By collecting data over extended periods (e.g., several weeks to months), it would be possible to detect rhythmic seizure patterns and identify individual cycles of seizure occurrence. This approach would allow for the prediction of seizures tailored to each individual, taking into account the variability in seizure cycles between patients [3].

Future studies should aim to further elucidate the mechanisms underlying these rhythmic patterns and their role in seizure generation. Investigating how these rhythms interact with other factors, such as seizure duration and inter-seizure intervals, could provide a more comprehensive understanding of seizure dynamics. Additionally, we plan to explore the neural mechanisms behind these cycles using computational models [10, 11, 31]. Computational modeling offers a powerful approach to simulate and analyze the complex interactions within neural networks that may drive these rhythmic patterns. By incorporating key physiological variables, such as neuronal excitability, synaptic connectivity, and network dynamics, these models will allow us to test various hypotheses about the emergence and stability of seizure cycles, potentially revealing new therapeutic targets[11]. Moreover, exploring the potential for modulating these rhythms through pharmacological or non-invasive interventions could open new avenues for epilepsy treatment.

## Conclusion

In conclusion, the analysis of seizure rhythms in this study using a TT rat model of epilepsy has revealed distinct and dynamic rhythmic patterns that vary in both period and power across individual rats. These findings contribute to our understanding of the temporal organization of seizures and suggest that rhythmic cycles play a significant role in the timing and predictability of seizure events. This knowledge could inform the development of targeted, timing-based interventions to improve epilepsy management.

The identification of multiday cycles in seizure occurrences provides significant insights into the temporal organization of seizures. These findings suggest that seizures are not entirely random but follow intrinsic rhythmic patterns, as evidenced in the TT rat model. Understanding these cycles has direct therapeutic implications, including the potential to forecast periods of heightened seizure likelihood. This predictive capability could allow for tailored interventions, such as adjusting medication dosages, scheduling neurostimulation therapies, or advising patients to take preventive measures during high-risk periods.

## Data availability statement

Data are available upon reasonable request from the corresponding authors. Code is available on Github (https://github.com/ParvinZE/Cycles-in-Epilepsy).

## Acknowledgments

The authors thank Dr Warwick Cheung for providing the rat recordings. This work was funded by the Australian Government under the Australian Research Council’s Training Centre in Cognitive Computing for Medical Technologies (project number ICI70200030).

